# Morph-specific investment in testes mass in a trimorphic beetle, *Proagoderus watanabei*

**DOI:** 10.1101/2021.05.09.443318

**Authors:** Jonathan M. Parrett, Eleanor M. Slade, Robert J. Knell

**Affiliations:** Evolutionary Biology Group, Faculty of Biology, Adam Mickiewicz University, Umultowska 89, 61–614, Poznań, Poland; School of Biological and Chemical Sciences, Queen Mary, University of London, Mile End Road, London E1 4NS, UK; Asian School of the Environment, Nanyang Technological University, 50 Nanyang Avenue, Singapore City 639798, Singapore

## Abstract

When competition between males for mates is intense it is common to find that some males will adopt alternative tactics for acquiring fertilisations, often involving the use of ‘sneak’ tactics whereby males avoid contests. These alternative tactics are sometimes associated with discrete differences in male morphology, with sneak males investing less in weaponry but more in traits such as testes which will give an advantage in sperm competition. In some cases it appears that males develop into more than two morphs, with a number of examples of tri- and even tetramorphic arthropod species being described. Here we analyse the scaling relations of the dung beetle species *Proagoderus watanabei*, which expresses two distinct weapon traits: paired head horns and a pronotal horn. We find that males of this species are trimorphic, with alpha males expressing long head horns and a pronotal horn, beta males with long head horns but no pronotal horn, and gamma males with short head horns only. We also find that alpha males invest less in testes than do beta or gamma males, indicating that beta and gamma males in this species probably experience higher risks of sperm competition than do alphas.

## Introduction

Strong intraspecific competition between males for mates is often associated with the use of alternative tactics by males of different status (Gross, 1996; Taborsky, Oliveira & Brockmann, 2008). In taxa as diverse as mites, fish, crickets, frogs, and bovids (Oliveira, Taborsky & Jane Brockmann, 2008) some males in a population will avoid competing directly with more dominant males, instead attempting to acquire matings while avoiding aggressive interactions. In many cases these alternative tactics are themselves associated with distinct morphological differences between males. In the insects these differences are common in the Coleoptera and especially in the superfamily Scarabaeoidea, where male dimorphism is frequently found in the Lucanidae (Matsumoto & Knell, 2017), the Dynastinae (McCullough *et al*., 2015), and the Scarabaeinae (Emlen *et al*., 2005; Simmons, Emlen & Tomkins, 2007). In these animals one morph, usually called the “major” morph typically invests heavily in weaponry such as horns or enlarged mandibles while the other, the “minor” morph bears reduced or no weaponry, leading to non-linear scaling relationships between weapon size and body size (Knell, 2009; McCullough *et al*., 2015). These dimorphic beetles are now regarded as important model systems for studies of development and sexual selection (Simpson, Sword & Lo, 2011; Casasa, Schwab & Moczek, 2017) and studies using them have provided considerable insight into patterns of resource allocation and the costs of investment in structures such as weapons and testes (Simmons & James Ridsdill-Smith, 2011).

In recent years, a number of examples of arthropods which appear to have more than two male morphs have been described. Three different male morphs have been found in some species of *Philotrypesis* fig wasp (Jousselin, van Noort & Greeff, 2004), a number of dung beetle species (Rowland & Emlen, 2009), stag beetles (Rowland & Emlen, 2009), (Iguchi, 2013; Matsumoto & Knell, 2017), a weta (Kelly & Adams, 2010), a weevil ((Rowland & Emlen, 2009) and two species of harvestman (Painting *et al*., 2015; Powell *et al*., 2020). In two lucanid species there is even evidence for four separate male morphs (Iguchi, 2013; Matsumoto & Knell, 2017).

Genetically based trimorphisms have been described from a number of lizard species (Sinervo & Lively, 1996; Sinervo *et al*., 2007), from one bird species (the Ruff, *Philomachus pugnax*, (*Küpper et al., 2016*), and the isopod *Paracerceis sculpta* (Shuster & Wade, 1991). The recently described arthropod trimorphisms are, however, likely to be conditional strategies rather than genetic polymorphisms (Rowland, Qualls & Buzatto, 2017), but to date we know little of their biology beyond the fact of their existence. The three morphs of male *Phylotrypesis* fig wasps appear from their morphology to have clear roles as large, aggressive male, small sneak male, and winged disperser ((Jousselin *et al*., 2004) but the roles played by the different morphs in other examples are less clear. (Painting *et al*., 2015) described morph specific behaviour during contests in the harvestman *Pantopsalis cheliferoides* but we know nothing further regarding questions like whether the different morphs in these animals behave differently, or whether they allocate resources in different ways during development in the way that many dimorphic arthropods following conditional strategies do.

One of the best studied consequences of alternative reproductive tactics was first pointed out by (Parker, 1990): When males adopt two different tactics, sneak and guard, then in general it is expected that the sneak males should invest more into traits associated with performance under sperm competition such as testes size (Kustra & Alonzo, 2020). This arises because the risks of sperm competition are greater for sneaks, who will usually be exposed to it whenever mating, than for guards, who will prevent other males from mating with females and so experience lower risks. The degree of difference in risk between sneaks and guards will itself depend on the frequency of sneaks in the population: when there are many sneaks the risk of sperm competition experienced by guards will be higher and so the differential between guards and sneaks is predicted to be lower (Parker, 1990; Gage, Stockley & Parker, 1995; Simmons *et al*., 2007). Testes size in species with alternative reproductive tactics has been studied in a variety of animals, especially fish (Kustra & Alonzo, 2020), but also in Onthophagine dung beetles (Simmons, Tomkins & Hunt, 1999; Simmons *et al*., 2007; Knell & Simmons, 2010) where these predictions have been found broadly to be supported.

Here, we analyse morphological data from males of a common Southeast Asian Onthophagine dung beetle, *Proagoderus watanabei* (*Ochi & Kon, 2002*). Both sexes of *P. watanabei* express paired head horns although these are considerably shorter even in the largest females than those of most males, many of whom bear striking long, curved horns. Both sexes also have considerable pronotal sculpting, which in some males develops into a single pronotal horn. In the original species description, (Ochi & Kon, 2002) described the males as occurring in three types, on the basis of the size of the head and pronotal horns but made no quantitative analysis of this. (Moczek, Brühl & Krell, 2004) analysed the allometric relationship between horn length and body size in both males and females of this species with a focus on comparing the scaling relationship of the head and pronotal horns but did not focus on the degree of polymorphism shown by the males. We extend this and find evidence to support a trimorphic model of male morphology in this species. We then analyse the relationship between testes mass and somatic mass in the different morphs to test whether patterns of investment in traits associated with sperm competition success vary between the different morphs.

## Methods

Individuals were sampled and measured as part of a community wide study (see (Parrett *et al*., 2019) for details). In brief, trapping was performed at the SAFE project ((Ewers *et al*., 2011) in Sabah, Malaysian Borneo in both 2011 and 2015. Pitfall traps baited with human dung were set across a habitat gradient, ranging from undisturbed and logged tropical forests to oil palm plantations. In 2015, live trapping was performed to gain measurements of testes mass, whereas, in 2011 beetles were killed during trapping and stored in ethanol. In 2015, beetles were housed in plastic containers with damp tissue paper prior to processing. All individuals were processed within 72 hrs of trapping. Individuals were killed by freezing, their total body mass taken, and then their testes were dissected out immediately and weighed using a Sartorius BP2215 balance. A calibration weight was used before each measurement. In all cases beetles were photographed from above and the side using a USB microscope and their pronotum width and horn length were measured using ImageJ v1.47 (Schneider, Rasband & Eliceiri, 2012).

### Statistical analysis

For the analysis of male morphology, we followed the procedure outlined in (Knell, 2009). After initial inspection of scatterplots of horn length vs. body size (pronotal width), we looked for evidence of bimodalism in frequency distributions of the ratios of horn length to body length. On the basis of potential morph allocations derived from these initial data explorations we analysed the scaling relationship of head horn length to body size in log-log space by comparing the AIC score for a series of candidate models with and without the various potential morph allocations as factors, plus a sigmoidal model as used in (Moczek *et al*., 2002). These data were collected from a number of replicate trapping stations so we used mixed effects models with replicate included as a random factor to control for the non-independence this introduces. We also fitted a breakpoint linear regression (Knell, 2009) without a random factor since methods for fitting mixed effects breakpoint models are not well developed.

Testes mass data was only available for beetles collected in 2015, and was analysed by fitting a series of candidate models with and without morph as an explanatory factor. The effect of size was controlled for by including somatic mass: because of multicollinearity issues we only used somatic mass as an explanatory variable and not pronotum width. Somatic mass was chosen because it is probably a more reliable indication of investment in body parts other than testes than is pronotum width. Initial exploratory analysis indicated the possibility of a curved relationship between testes mass and somatic mass so a quadratic term was included in one candidate model.

All analyses were carried out in R v.4.03 (R Development Core Team, 2021) and mixed effects models were fitted using the lme4 package (Bates *et al*., 2015). The breakpoint regression was fitted using the Segmented package (Muggeo & Muggeo, 2017). Full code and results for the analysis and data visualisation are included in the supplementary material.

## Results

### Analysis of male morphology

The relationship between head horn length and body size is non-linear and the histogram for the ratio of head horn length to pronotum width is bimodal with a minimum between the two peaks at 0.42 (Figure 1). Many beetles (45%) have no pronotal horn and there appears to be a qualitative difference between those with and those without horns: there is only one animal in the dataset with a pronotal horn which is less than 1mm long. The histogram for the ratio of pronotal horn length to pronotum width is unimodal when those animals with no pronotal horn at all are excluded. On this basis we can divide these male beetles into three groups (figure 2): beetles with small head horns and no pronotal horn (gamma morphs), beetles with large head horns and no pronotal horn (beta morphs) and beetles with large head horns and a pronotal horn (alpha morphs).

**Figure 1.**
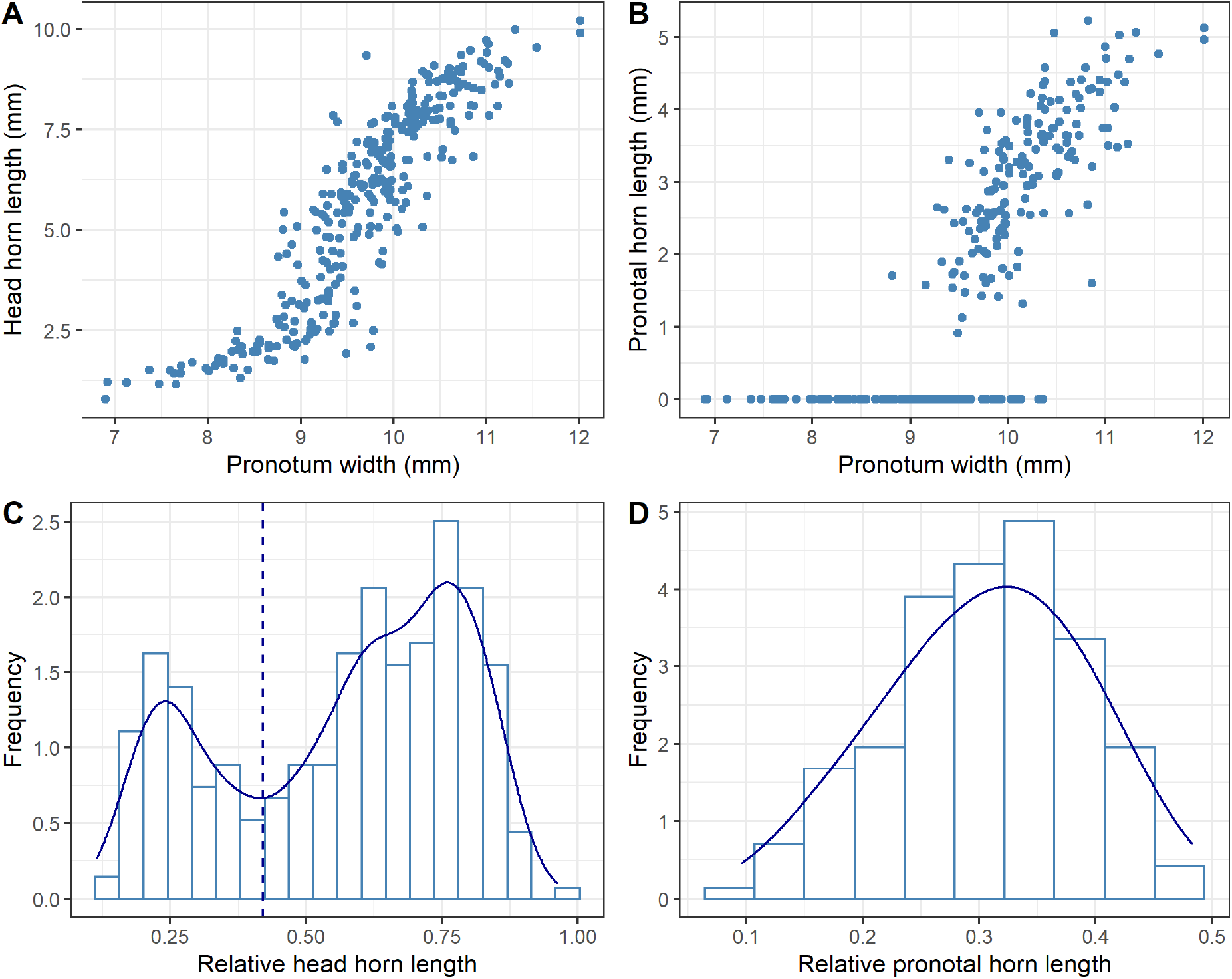
Horn lengths and ratios of horn length to body size (pronotum width). A: head horn length plotted against pronotum width. B: Pronotal horn length against pronotum width. C: Frequency distribution of the ratio of head horn length to pronotum width. The solid line shows a kernel density estimator and the vertical dashed line shows the minimum between the two peaks at a ratio of 0.42. D. Frequency histogram of the ratio of pronotal horn length to pronotum width, with a kernel density estimator shown as for C.

**Figure 2.**
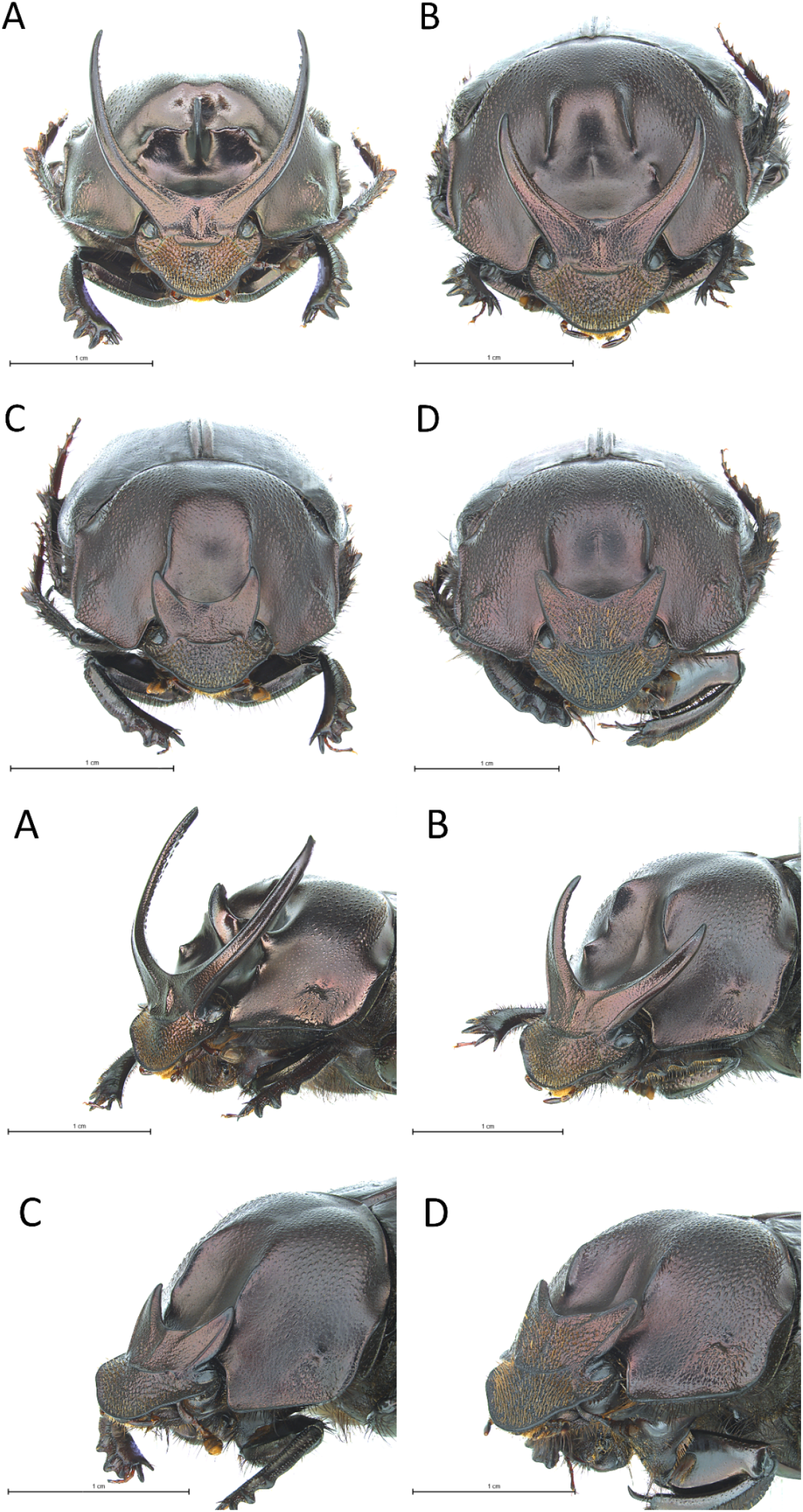
Frontal (top four panels) and angled (lower four panels) views of A: an alpha male morph, B: a beta male morph, C: a gamma male morph and D: a female of P. watanabei. Image credit Xin Rui Ong, TEE Lab, NTU Singapore.

Of our candidate set of models, the model with a discontinuous relationship split into the three morphs outlined above has by far the lowest AIC score and is therefore our preferred model for *P. watanabei* — note that including morph allocation based on the pronotal horn makes a considerable improvement to the model’s goodness of fit when explaining the patterns in the head horns (Figure 3, Table 1).

**Figure 3.**
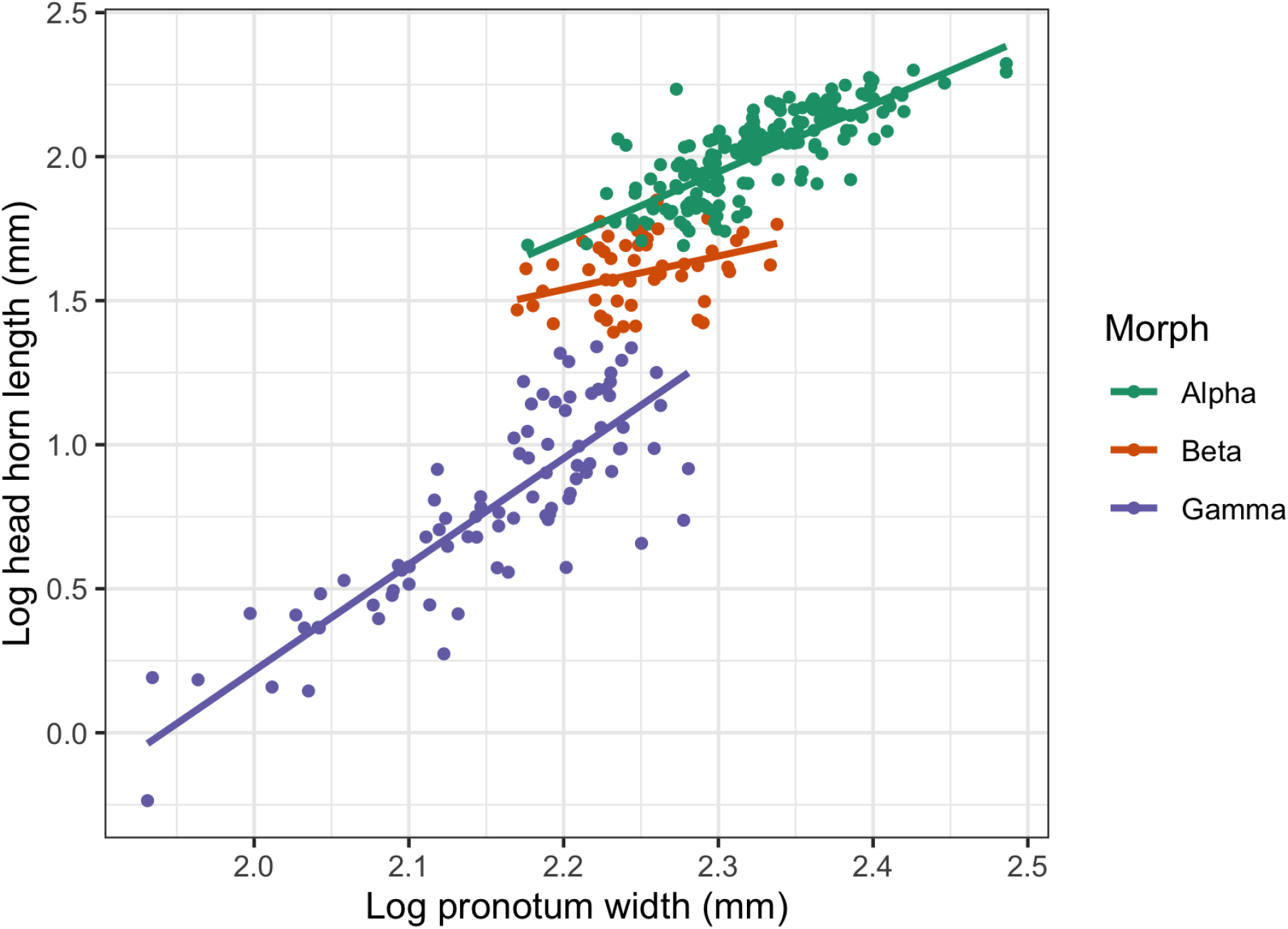
Scaling relationship between head horn length and pronotum width plotted on a log-log scale. Colours indicate morph allocations. Lines show the predicted values from the fixed effects in the fitted statistical model.

**Table 1.** Candidate models to describe the scaling relationship between log head horn length and log pronotum width and their AIC and ΔAIC scores. All models except M3 were fitted as mixed effects models with replicate as a random factor.

| Candidate Model | AIC | ΔAIC |
| --- | --- | --- |
| M1: simple linear regression | -12.0 | 368.9 |
| M2: polynomial regression with a quadratic term | -19.48 | 361.4 |
| M3: breakpoint linear regression fitted with the Segmented R package (Muggeo & Muggeo, 2017) | -6.56 | 374.3 |
| M4: four-parameter non-linear sigmoid model as used by (Moczek *et al.*, 2002) | -102.41 | 278.5 |
| M5: linear model with “morph” as a factor and pronotum width as a continuous predictor variable, where “morph” is a two-level factor based on head horn length. | -291.7 | 89.2 |
| M6: linear model with “morph” as a factor and pronotum width as a continuous predictor variable, where “morph” is a two-level factor based on the presence or absence of a pronotal horn. | -153.1 | 227.8 |
| M7: linear model with “morph” as a factor and pronotum width as a continuous predictor variable, where “morph” is a three-level factor dividing the males into three morphs on the basis of both head horn length and the presence or absence of a pronotal horn. | -380.9 | 0 |

### Analysis of testes mass

Out of our candidate set of models for testes mass, the AIC scores indicate the strongest support for the model with the quadratic term (T4). Models without this term (T2 and T3) have weak support with a delta AIC of about 6.6 in both cases (Table 2). The nesting rule (Harrison *et al*., 2017) suggests that we should discount model T3, so we conclude that there is strong support for an effect of both morph and somatic mass on testes mass, slightly weaker support for a quadratic effect of somatic mass and little support for an interaction between morph and somatic mass. Examining the coefficients table for the best model T4 (see Supplementary information) tells us that alpha morphs are investing less in testes mass than both beta and gamma morphs but there appears to be little difference in testes mass between the beta and gamma morphs (Figure 4), so beta and gamma morphs had testes that were roughly 8-10mg heavier than those of alpha morphs when the effect of somatic mass was accounted for.

**Table 2.** Candidate models to describe the relationship between testes mass, somatic mass and morph, with their AIC and ΔAIC scores.

| Candidate Model | AIC | ΔAIC |
| --- | --- | --- |
| T1: testes mass explained by somatic mass only | -1348.9 | 13.6 |
| T2: testes mass explained by main effects of morph and somatic mass | -1356.0 | 6.58 |
| T3: testes mass explained by main effects and interaction of morph and somatic mass | -1355.9 | 6.65 |
| T4: testes mass explained by main effects of morph and somatic mass plus a quadratic term for somatic mass. | -1362.6 | 0 |

**Figure 4.**
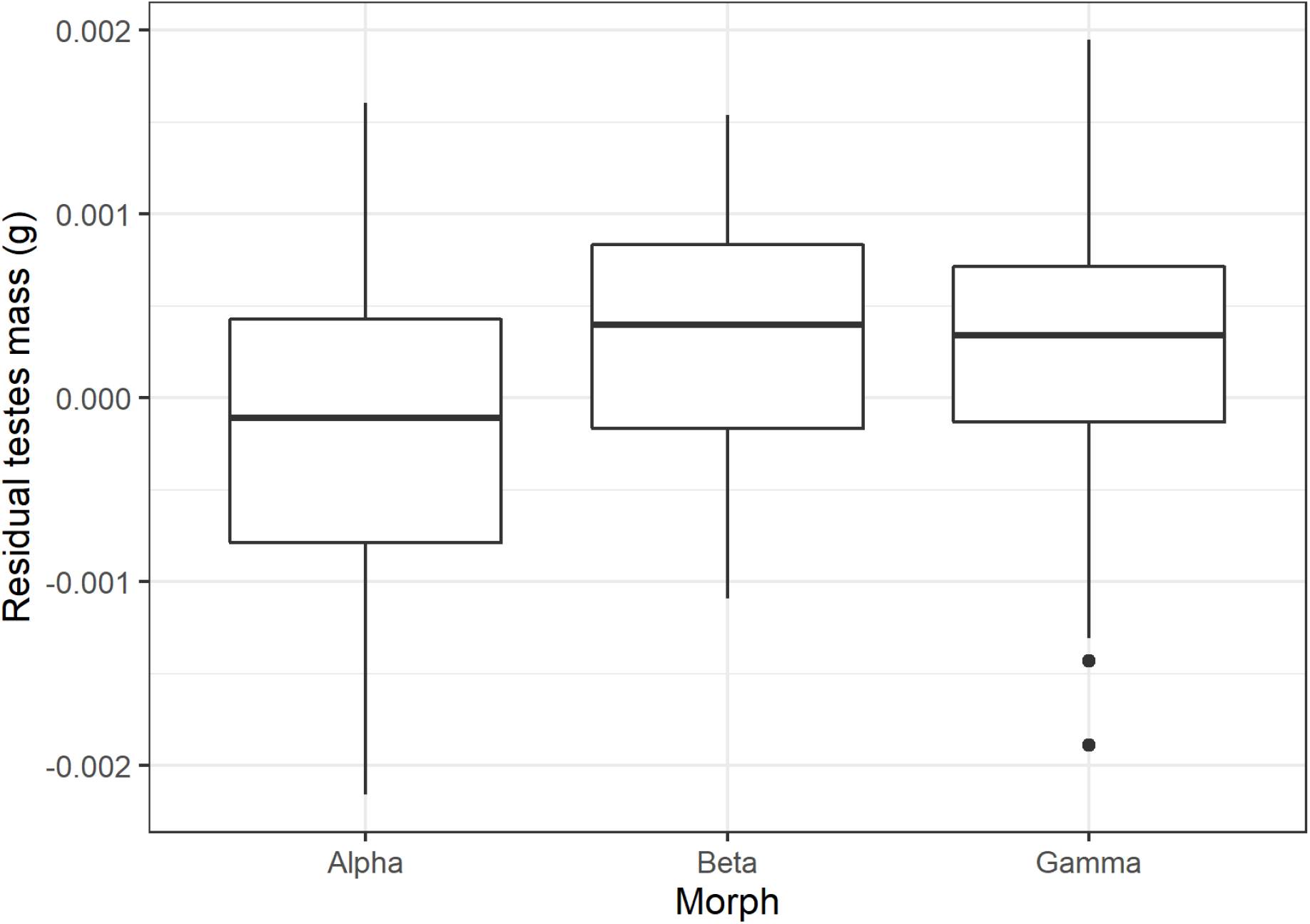
The effect of morph on testes mass. The boxplots show the residuals from a mixed effects model with main effects of somatic mass and somatic mass squared plotted against morph. The alpha morphs tend to have smaller testes when compared to the beta and gamma morphs.

## Discussion

On the basis of the scaling relationships of weapons and body size we find that *P. watanabei* males group into three separate morphs, and that those males with large head horns and pronotal horns, the alpha morph, invest relatively less in testes mass than do either the beta males (large head horns but no pronotal horn) or the gamma males (small head horns, no pronotal horn). Previous studies of trimorphic arthropod males have analysed differences in the scaling relationships of single weapon traits only (*Rowland & Emlen, 2009; Painting *et al*., 2015; Matsumoto & Knell, 2017; Powell et al*., 2020). The males of many beetle species carry horns on both their heads and their pronota (Emlen *et al*., 2005; Emlen, 2008; Knell, 2011), and we suggest that further study of some of these might reveal similar complex polymorphisms to the one described here.

Our finding of complex scaling relations for the head horns and trimorphic males in *P. watanabei* contrasts with the previous investigation of male horn scaling in *P. watanabei* which concluded that the scaling relationship for the head horns was best described by a linear model and that males are essentially monomorphic (Moczek et al. 2004). There are also similarities between both datasets: for example, Moczek et al (2004) also noted a clear binary difference between males which do or do not express prothoracic horns and that the allometry of head horns was best described by a non-linear relationship. By using our considerably larger dataset (304 compared to 71) we were able to detect a discontinuous relationship between head horn length and pronotum width, and by assigning a binary value for prothoracic horn expression we were able to allocate males to a third ‘intermediate’ morph (i.e. beta males), confirming the qualitative description of three male morphs in this beetle suggested by Ochi & Kon (2002).

The aggressive tactics adopted by large, well armed beetles and the sneak tactics used by small, unarmed males are well known from dimorphic species and it is likely that the alpha and gamma males of *P. watanabei* behave in similar ways. But what of the beta males? One possibility is that they might adjust their reproductive behaviours plastically depending on context: using their horns and adopting aggressive behaviours to monopolise females in the presence of gamma males, and using sneak tactics in the presence of alpha males. The function of the prothoracic horn is not known but based on the pronotum shape of males which do not express a prothoracic horn (figure 2) it seems likely that it acts as a signal to other males, conveying information on overall body size. Previous work on scarab species with dimorphic males has found that major males tend to have smaller testes for their body size than minor males (Simmons *et al*., 1999, 2007). This is seen as an adaptation to different degrees of risk of sperm competition between these morphs, and it is likely that the patterns found here reflect this as well. It seems, therefore, that in this species the alpha males are at reduced risk of sperm competition than the beta and gamma males. On this basis we can speculate that alpha males, as with major males in dimorphic species, are able to defend females against most rivals thereby reducing the risk of sperm competition. beta males of *P. watanabei* invest in traits that function in both physical contests (horns) and in sperm competition (testes). If they are less able to defend females than alphas then their risk of sperm competition will be increased since they will more often be ousted by rival males who will then mate with the female that they were defending. The gamma males, with reduced horns, are likely to be sneak mating specialists and so will also experience high levels of sperm competition. Why there is no difference between beta and gamma males is not currently clear: it is possible that overall there is no difference because the balance of costs (resources used) and benefits (better outcomes when exposed to sperm competition) is the same for both morphs. The alternative is that because of a type II error (i.e. a ‘false negative’) we have failed to detect an existing small difference. There is obvious potential here for further research to determine what behavioural differences exist between male morphs and to describe the function of different horn types in *P. watanabei*..

## Acknowledgements

We are grateful to the Land use Options for Maintaining BiOdiversity and eKosystem functions (LOMBOK) Project research assistants for their assistance in the field. We thank Marx Lim and Xin Rui Ong, TEE lab., for the photos in Figure 2. Arthur Chung, Glen Reynolds and SEARRP, The SAFE Project, The Maliau Basin Management Committee, and the Sabah Biodiversity Council, Malaysia, for their support in obtaining permits and research permissions. Samples were collected under SaBC access licence number JKM/MBS.1000-2/2(381) to EMS and JMP. EMS was supported by a UK Natural Environment Research Council grant (NE/K016407/1), and NTU start-up grant. JMP was supported by a QMUL postgraduate studentship and funding from QMPGRF.

## Data sharing statement

Data from this study are archived on the SAFE project data repository at https://zenodo.org/record/3342495 (DOI10.5281/zenodo.3342495). Code for the extraction of the *P. watanabei* data from this large dataset and all the analysis carried out for this manuscript is included in the supplementary material.

