## Supplementary material for "Morph-specific investment in testes mass in a trimorphic beetle, *Proagoderus watanabei*": Proagoderus_Analysis_April_2021.html


Code 

- Show All Code
- Hide All Code

### Supplementary material for Morph-specific investment in testes mass in a trimorphic beetle, *Proagoderus watanabei*

###### Rob Knell

###### 28 February 2021

Authors: Jonathan M. Parrett^1,2^, Eleanor M. Slade^3^ & Robert J. Knell^2,4^

1 Evolutionary Biology Group, Faculty of Biology, Adam Mickiewicz University, Umultowska 89, 61–614, Poznań, Poland 2 School of Biological and Chemical Sciences, Queen Mary, University of London, Mile End Road, London E1 4NS, UK 3 Asian School of the Environment, Nanyang Technological University, 50 Nanyang Avenue, Singapore City 639798, Singapore 4

#### Setup

Load packages

```
library(segmented)
library(ggplot2)
library(lme4)
library(lmerTest)
library(cowplot)
library(readxl)
library(curl)
library(dplyr)
```

Imports dataset from SAFE repository, extracts data for male *P. watanabei* only, ensures numeric variables are numeric and factors are factors.

```
# Import spreadsheet
url <- "https://zenodo.org/record/3342495/files/SAFE_database_dung_beetle_sexual_selection_data_final_uploaded.xlsx"
destfile <- "SAFE_database_dung_beetle_sexual_selection_data_final_uploaded.xlsx"
curl_download(url, destfile)
dat1 <- read_excel(destfile,
                                     sheet = "morphology_data",
                                     range = "a7:m7748")

# Extract data we want
males <- dat1 %>% filter(species == "Proagoderus_watanabei" & sex == "male")

# readxl has issues with variable classes when NAs present, this sorts it out
males$horn_front <- as.numeric(males$horn_front)
males$horn_side <- as.numeric(males$horn_side)
males$pronotum_horn_front <- as.numeric(males$pronotum_horn_front)
males$pronotum_horn_side <- as.numeric(males$pronotum_horn_side)
males$total_mass <- as.numeric(males$total_mass)
males$testes_mass <- as.numeric(males$testes_mass)

# adjmass = somatic mass
males$adjmass <- males$total_mass - males$testes_mass

males$sex <- as.factor(males$sex)
males$year <- as.factor(males$year)

rm(url, destfile, dat1)
```

#### Male polymorphism

In order to investigate the possibility of multiple male morphs with different horn morphologies we followed the procedure suggested in Knell (2009)^1^. Following initial data visualisation candidate morphs were identified from histograms of the ratio of horn length to body size (pronotum width in this case) and a series of candidate models compared via their AIC scores to determine the morph allocation and model structure which best explains the relationship between horn length (in this case head horn length) and body size.

##### Head horn to pronotum width plot

```
p1 <- ggplot(data = males) +
    aes(x = pronotum_width, y = horn_side) +
    geom_point(colour = "steelblue") +
    theme_bw() +
    theme(panel.grid.minor = element_blank()) +
    theme(panel.grid.major = element_line(size = 0.2)) +
    labs(x = "Pronotum width (mm)", y = "Head horn length (mm)")
p1
```

**Figure 1. Head horn length plotted against pronotum width.**

The relationship between head horn length and body size appears distinctly non-linear, indicating possible polymorphism.

##### Pronotal horn to pronotum width plot

```
p2 <- ggplot(data = males) +
    aes(x = pronotum_width, y = pronotum_horn_side) +
    geom_point(colour = "steelblue") +
    theme_bw() +
    theme(panel.grid.minor = element_blank()) +
    theme(panel.grid.major = element_line(size = 0.2)) +
    labs(x = "Pronotum width (mm)", y = "Pronotal horn length (mm)")
p2
```

**Figure 2. Pronotal horn length plotted against pronotum width.**

Pronotal horns appears to fall into two distinct groups. One group of males have no pronotal horn at all, and then a second group of males have horns, with no males apparently bearing very small horns. This also indicates potential polymorphism between those males with and without pronotal horns. There might be further structure in the horn lengths of the males with pronotal horns but this can’t be seen in this scatterplot.

##### Histogram of head horn length to pronotum width ratio

```
# Calculate relative horn lengths

males$RH <- males$horn_side/males$pronotum_width
males$sec_RH <- males$pronotum_horn_side/males$pronotum_width


# Plot histogram 
p3 <- ggplot(data = males) +
    aes(RH) +
    geom_histogram(aes(y = ..density..),
                                 fill = "white",
                                 colour = "steelblue",
                                 bins = 20) +
    geom_density(
        lty = 1,
        size = 0.5,
        col = "darkblue",
        bw = 0.05
    ) +
    geom_vline(
        xintercept = 0.42,
        linetype = 2,
        color = "darkblue",
        size = 0.5
    ) +
    theme_bw() +
    theme(panel.grid.minor = element_blank()) +
    theme(panel.grid.major = element_line(size = 0.2)) +
    labs(x = "Relative head horn length", y = "Frequency")
p3
```

**Figure 3. Histogram of the ratios of head horn length to pronotum width**, showing a kernel density estimator. The vertical line shows the cutoff between morphs at a ratio of 0.42

This histogram of the ratios of head horn length to pronotum width shows two distinct peaks: one group of males with relatively small horns and one with relatively large horns.

```
# Plot histogram of pronotal horn length to pronotum width ratio
# Not plotting males with no pronotal horn
p4 <- ggplot(data = males[males$pronotum_horn_side > 0,]) +
    aes(sec_RH) +
    geom_histogram(aes(y = ..density..),
                                 colour = "steelblue",
                                 fill = "white",
                                 bins = 10) +
    geom_density(
        lty = 1,
        size = 0.5,
        col = "darkblue",
        bw = 0.05
    ) +
    theme_bw() +
    theme(panel.grid.minor = element_blank()) +
    theme(panel.grid.major = element_line(size = 0.2)) +
    labs(x = "Relative pronotal horn length", y = "Frequency")
p4
```

**Figure 5. Histogram of the ratios of pronotal horn length to pronotum width** showing a kernel density estimator. Individuals with a pronotal horn length of zero are not shown.

The histogram of the ratios of pronotal horn length to pronotum width shows only a single peak with no indication of any further structure when the males with no pronotal horn are excluded.

```
# Assemble plots into paper figure 1
# fig_1 <- plot_grid(p1, p2, p3, p4, labels = c("A", "B", "C", "D"))
# 
# ggsave("fig_1.png", plot = fig_1, device = "png", width = 20, height = 16, units = "cm")
```

#### Morph allocation

Based on the plots above we can allocate males to morphs in two ways. Firstly, we can divide males on the basis of the ratio of head horn to pronotum width, with the divide at 0.42. Secondly, we can divide males into those with and those without a pronotal horn.

This gives three morphs in total: those with large head horns and a pronotal horn (alphas), those with large head horns but no pronotal horn (betas) and those with small head horns and no pronotal horn (gammas). There are no males with small head horns and a pronotal horn.

```
males$head.morph<-cut(males$RH,c(-0.1,0.42,1.3),labels=c("minor","major"))
males$pron.morph<-cut(males$sec_RH,c(-0.1,0.05,1.3),labels=c("no","yes"))

males$Morph <- ifelse(males$head.morph == "major" & males$pron.morph == "yes", "Alpha", ifelse(males$head.morph == "major" & males$pron.morph == "no", "Beta", "Gamma"))

males$Morph <- as.factor(males$Morph)
```

##### Fit candidate models

To determine which models provide the best fit to the data for log head horn length and log pronotum width we fit a series of candidate models and compare their AIC scores. The models are all mixed effects models with replicate as a random factor aside from the breakpoint regression since we are not aware of methods for including random factors in breakpoint regression.

m1: simple linear regression

m2: second order polynomial

m3: breakpoint linear regression

m4: sigmoidal model

m5: discontinuous relationship described by two morphs based on head horn length

m6: discontinuous relationship described by two morphs based on pronotal horn length

m7: discontinuous relationship described by three morphs

```
# Simple linear regression
m1<-lmer(log(horn_side)~log(pronotum_width) + (1|replicate), REML = FALSE, data = males)

# Second order polynomial
m2<-lmer(log(horn_side)~log(pronotum_width)+I(log(pronotum_width)^2) + (1|replicate), REML = FALSE, data = males)

# Breakpoint model fitted using segmented package
y<-log(males$horn_side)
x<-log(males$pronotum_width)
c<-lm(y~x)
m3<-segmented(c,seg.Z=~x,psi=2.2)

# Sigmoidal model fitted using non-linear regression

sig_mod <- ~(y + (a * X ^ b)) / (c ^ b + X ^ b)


sig_mod_gradient <- deriv(
    sig_mod, 
    namevec = c("y", "a", "b", "c"), 
    function.arg=c("X","y", "a", "b", "c")
)

m4 <-
  nlmer(log(horn_side) ~ sig_mod_gradient(y,a,b,c,X = log(pronotum_width)) ~
          (c|replicate), 
  data = males,
  start = c(
    y = 0,
    a = 0.6,
    b = 30,
    c = 2.2)
  )

# Model with discontinuous relationship with two morphs (large and small head horns)
m5<-lmer(log(horn_side)~log(pronotum_width)*head.morph + (1|replicate), REML = FALSE, data = males)

# Model with discontinuous relationship with two morphs (pronotal horn present/absent)
m6<-lmer(log(horn_side)~log(pronotum_width)*pron.morph + (1|replicate), REML = FALSE, data = males)

# Model with discontinuous relationship with three morphs
m7<-lmer(log(horn_side)~log(pronotum_width)*Morph + (1|replicate), REML = FALSE, data = males)


temp <- AIC(m1, m2, m3, m4, m5, m6, m7)

temp$deltaAIC <- round(temp$AIC - min(temp$AIC), 2)

temp$AIC <- round(temp$AIC, 2)

temp
```

```
##    df     AIC deltaAIC
## m1  4  -11.99   368.89
## m2  5  -19.48   361.41
## m3  5   -6.56   374.33
## m4  6 -102.41   278.48
## m5  6 -291.68    89.21
## m6  6 -153.12   227.77
## m7  8 -380.89     0.00
```

m7, the model with three morphs, has by far the lowest AIC score. This indicates that it is the best at explaining the relationship between log head horn length and log pronotum width.

##### Diagnostics

```
plot(m7)
```

**Figure 6. Residuals versus fitted values for the three-morph model relating log head horn length to log pronotum width.**

No indication of any problems from the diagnostic plot.

##### Model summary

```
summary(m7)
```

```
## Linear mixed model fit by maximum likelihood . t-tests use Satterthwaite's
##   method [lmerModLmerTest]
## Formula: log(horn_side) ~ log(pronotum_width) * Morph + (1 | replicate)
##    Data: males
## 
##      AIC      BIC   logLik deviance df.resid 
##   -380.9   -351.2    198.4   -396.9      296 
## 
## Scaled residuals: 
##     Min      1Q  Median      3Q     Max 
## -3.3840 -0.5901  0.0190  0.5857  3.1057 
## 
## Random effects:
##  Groups    Name        Variance Std.Dev.
##  replicate (Intercept) 0.001372 0.03705 
##  Residual              0.015184 0.12323 
## Number of obs: 304, groups:  replicate, 12
## 
## Fixed effects:
##                                Estimate Std. Error       df t value Pr(>|t|)
## (Intercept)                     -3.4491     0.4361 301.5799  -7.909 4.96e-14
## log(pronotum_width)              2.3459     0.1876 301.3813  12.507  < 2e-16
## MorphBeta                        2.4287     1.0986 300.0436   2.211   0.0278
## MorphGamma                      -3.7014     0.5710 298.1434  -6.482 3.74e-10
## log(pronotum_width):MorphBeta   -1.1830     0.4863 300.3003  -2.433   0.0156
## log(pronotum_width):MorphGamma   1.3368     0.2541 298.3576   5.261 2.75e-07
##                                   
## (Intercept)                    ***
## log(pronotum_width)            ***
## MorphBeta                      *  
## MorphGamma                     ***
## log(pronotum_width):MorphBeta  *  
## log(pronotum_width):MorphGamma ***
## ---
## Signif. codes:  0 '***' 0.001 '**' 0.01 '*' 0.05 '.' 0.1 ' ' 1
## 
## Correlation of Fixed Effects:
##             (Intr) lg(p_) MrphBt MrphGm l(_):MB
## lg(prntm_w) -0.999                             
## MorphBeta   -0.357  0.358                      
## MorphGamma  -0.745  0.746  0.299               
## lg(prn_):MB  0.345 -0.346 -1.000 -0.290        
## lg(prn_):MG  0.719 -0.720 -0.292 -0.999  0.284
```

##### Visualisation

```
# Calculate predicted values from fixed effects in mixed effects model

pred <-
  ifelse(
    males$Morph == "Alpha",
    -3.449 + 2.346 * log(males$pronotum_width),
    ifelse(
      males$Morph == "Beta",
      -3.449 + 2.429 + (2.346 - 1.183) * log(males$pronotum_width),
      -3.449 - 3.701 + (2.346 + 1.337) * log(males$pronotum_width)
    )
  )

p6 <- ggplot(data = males) +
  aes(x = log(pronotum_width), y = log(horn_side), colour = Morph) +
    geom_point(shape = 16) +
  geom_line(aes(y = pred), size = 1) +
    #geom_smooth(method = "lm", se = FALSE) +
    theme_bw() +
    theme(panel.grid.minor = element_blank()) +
    theme(panel.grid.major = element_line(size = 0.2)) +
    scale_color_brewer(palette = "Dark2") +
    labs(x = "Log pronotum width (mm)", y = "Log head horn length (mm)")
p6
```

**Figure 7. Log head horn length plotted against log pronotum width.** Colours indicate morph and lines show the predicted values from the fixed effects in the fitted three morph model.

```
# ggsave("fig_2.png", plot = p6, device = "png", width = 14, height =10, units = "cm")
```

#### Analysis of testes mass

Testes mass data is only available for beetles collected in 2015. We have two potential variables for overall body size, namely pronotum width and somatic body mass. Strong multicollinearity means that we will only use the somatic mass as a predictor variable for body size, not pronotum width as well. Somatic mass is used here since it is probably a more reliable indication of investment in body parts other than testes than is pronotum width. Initial exploratory analysis indicated the possibility of a curved relationship between testes mass and somatic mass so a quadratic term was included in one candidate model.

These data were collected from a number of replicate trapping stations and in order to control for the non-independence this introduces we used random intercept mixed effects models with replicate as the random factor. Models are:

x1: testes mass explained by somatic mass only

x2: testes mass explained by main effects of morph and somatic mass

x3: testes mass explained by main effects and interaction of morph and somatic mass

x4: testes mass explained by main effects of morph and somatic mass plus a quadratic term for somatic mass.

```
x1<-lmer(testes_mass ~ 
                    adjmass + 
                    (1|replicate),
                  data=males)

x2<-lmer(testes_mass~Morph + 
                    adjmass +
                    (1|replicate),
                  data=males)

x3<-lmer(testes_mass~Morph * 
                    adjmass +
                    (1|replicate),
                  data=males)

x4<-lmer(testes_mass ~  adjmass + 
                    I(adjmass^2) + 
                    Morph +
                    (1|replicate),
                  data=males)

temp <- anova(x1, x2, x3, x4)

AICs <- data.frame(model = rownames(temp), AIC = round(temp$AIC,1), deltaAIC =  round(temp$AIC - min(temp$AIC), 2))

AICs
```

```
##   model     AIC deltaAIC
## 1    x1 -1348.9    13.63
## 2    x2 -1356.0     6.58
## 3    x4 -1362.6     0.00
## 4    x3 -1355.9     6.65
```

The AIC scores indicate the strongest support for model x4, with the quadratic term. models x2 and x3 have weak support with a delta AIC of about 6.6 in both cases. The nesting rule (Harrison 2017^2^) suggests that we should discount model x3 so we conclude that there is strong support for an effect of both morph and somatic mass on testes mass, slightly weaker support for a quadratic effect of somatic mass and little support for an interaction between morph and somatic mass.

##### Model summary

```
summary(x4)
```

```
## Linear mixed model fit by REML. t-tests use Satterthwaite's method [
## lmerModLmerTest]
## Formula: testes_mass ~ adjmass + I(adjmass^2) + Morph + (1 | replicate)
##    Data: males
## 
## REML criterion at convergence: -1310.7
## 
## Scaled residuals: 
##     Min      1Q  Median      3Q     Max 
## -2.5602 -0.5865  0.1429  0.6753  2.3901 
## 
## Random effects:
##  Groups    Name        Variance  Std.Dev. 
##  replicate (Intercept) 1.424e-07 0.0003774
##  Residual              6.904e-07 0.0008309
## Number of obs: 122, groups:  replicate, 12
## 
## Fixed effects:
##                Estimate Std. Error         df t value Pr(>|t|)    
## (Intercept)  -2.054e-03  1.477e-03  1.159e+02  -1.391 0.166987    
## adjmass       2.718e-02  6.864e-03  1.146e+02   3.960 0.000131 ***
## I(adjmass^2) -2.443e-02  8.021e-03  1.137e+02  -3.046 0.002881 ** 
## MorphBeta     8.353e-04  2.492e-04  1.137e+02   3.352 0.001088 ** 
## MorphGamma    9.664e-04  2.569e-04  1.169e+02   3.762 0.000265 ***
## ---
## Signif. codes:  0 '***' 0.001 '**' 0.01 '*' 0.05 '.' 0.1 ' ' 1
## 
## Correlation of Fixed Effects:
##             (Intr) adjmss I(d^2) MrphBt
## adjmass     -0.978                     
## I(adjmss^2)  0.932 -0.985              
## MorphBeta   -0.290  0.210 -0.151       
## MorphGamma  -0.521  0.400 -0.299  0.418
```

From the coefficients table it’s clear that both Beta and Gamma males have greater relative investment in testes than do Alphas.

##### Visualisation

```
# generate residuals from a model without morph as an explanatory factor

resids <- residuals(
              lmer(testes_mass ~ 
                       adjmass + 
                       I(adjmass^2) +
                       (1|replicate),
                     data=males))


# Plot boxplot of residual testes mass once somatic mass has been partialled out versus morph

p7 <- ggplot(data = data.frame(resids, Morph = males$Morph[males$year == "2015"])) +
  aes(x = Morph, y = resids) +
    geom_boxplot() +
    theme_bw() +
    theme(panel.grid.minor = element_blank()) +
    theme(panel.grid.major = element_line(size = 0.2)) +
    labs(x = "Morph", y = "Residual testes mass (g)")

p7
```

**Figure 8. Boxplot of residual testes mass once somatic mass has been controlled for plotted against morph.**

```
# ggsave("fig_3.png", plot = p7, device = "png", width = 14, height =10, units = "cm")
```

---

1. Knell, R.J. (2009) On the analysis of non-linear allometries. *Ecological entomology*, **34**, 1–11.↩︎
2. Harrison, X.A., Donaldson, L., Correa-Cano, M.E., Evans, J., Fisher, D.N., Goodwin, C., Robinson, B., Hodgson, D.J. & Inger, R. (2017) Best practice in mixed effects modelling and multi-model inference in ecology. *PeerJ*, **6**, e4794.↩︎
